# MACMIC Reveals Dual Role of CTCF in Epigenetic Regulation of Cell Identity Genes

**DOI:** 10.1101/2020.08.12.247361

**Authors:** Guangyu Wang, Bo Xia, Man Zhou, Jie Lv, Dongyu Zhao, Yanqiang Li, Yiwen Bu, Xin Wang, John P. Cooke, Qi Cao, Min Gyu Lee, Lili Zhang, Kaifu Chen

**Author notes:** Corresponding: Kaifu Chen. These authors contributed equally (Guangyu Wang, Bo Xia).

## Abstract

Numerous studies of relationship between epigenomic features have focused on their strong correlation across the genome, likely because such relationship can be easily identified by many established methods for correlation analysis. However, two features with little correlation may still colocalize at many genomic sites to implement important functions. There is no bioinformatic tool for researchers to specifically identify such feature pair. Here, we develop a method to identify feature pair in which two features have maximal colocalization but minimal correlation (MACMIC) across the genome. By MACMIC analysis of 3,385 feature pairs in 15 cell types, we reveal a dual role of CTCF in epigenetic regulation of cell identity genes. Although super-enhancers are associated with activation of target genes, only a subset of super-enhancers colocalized with CTCF regulate cell identity genes. At super-enhancers colocalized with CTCF, the CTCF is required for the active marker H3K27ac in cell type requiring the activation, and also required for the repressive marker H3K27me3 in other cell types requiring the repression. Our work demonstrates the biological utility of the MACMIC analysis and reveals a key role for CTCF in epigenetic regulation of cell identity.

## INTRODUCTION

As DNA sequencing data expands at an unprecedented speed, genomic (including epigenomic) data such as RNA-seq, ChIP-seq and genome sequencing data can be conveniently collected from public databases. Each set of sequencing data is typically collected to investigate a genomic (including epigenomic) feature across the genome, e.g., RNA-Seq dataset to investigate the expression profile of all genes in a genome, ChIP-Seq dataset to investigate a histone modification or the binding of a transcription factor at individual sites across the genome. It is commonly recognized that the function of a genome cannot be fully understood by studying a single genomic feature. Many studies showed that analysis of correlation between two genomic features had a strong potential to identify their regulatory relationship in an important biological process^1^. For instance, a strong positive correlation between the binding intensity of a protein near individual genes and the expression level of these genes might help define the protein to be an activator of transcription^2^. By focusing on the correlation between the RNA expression and a histone modification, the roles of individual histone modifications in the activation or repression of transcription have also been recognized^3,4^.

However, in many aspects of informatics, the representation of knowledge can be more efficient by using a combination of uncorrelated features^5^. In other words, highly correlated features often contain redundant information^6^. For example, whereas the dozens of pluripotent factors such as Oct4, Sox2, Klf-4, and c-Myc, are all useful to predict genes expressed in stem cells ^7-9^, combining some pluripotent factors with endothelial lineage factors such as Lmo2 and Erg would add power to also predict genes expressed in endothelial cells; therefore, it can be more powerful using combined information from transcription factors with distinct functions, as opposed to an analysis using the transcription factors with similar effects on a shared set of target genes. More importantly, colocalization of low-correlation chromatin features may still happen in a biologically considerable manner to implement important functions. For instance, the histone modifications H3K27me3 and H3K4me3 are known to be associated with repression and activation of transcription in differentiated cells, respectively^10^. As a result, they show negative correlation and often occur at different genes in somatic cell types^11^. However, these two markers lose the correlation and colocalize at a large set of genes in embryonic stem cells^12^. It is well known now that the colocalizations of H3K27me3 and H3K4me3 in embryonic stem cells define bivalent chromatin domains, which are functionally distinct from both the repressive domains associated with H3K27me3 and the active domains associated with H3K4me3. Theses bivalent chromatin domains play a unique role in embryonic stem cells to maintain a bivalent status of the lineage factors for individual somatic cell types ^13-15^. Therefore, analyzing colocalization of two chromatin features with globally low correlation in a cell has the potential to reveal novel biological mechanisms. However, little is known yet about the biological implications of such colocalization for the other chromatin features beyond H3K4me3 and H3K27me3. Therefore, the community is in need of a robust method to identify and understand the biologically important colocalization of uncorrelated chromatin features in a cell.

In this study, we utilized mutual information^16^ as an indication for general correlation (relevance) between a pair of genomic features, and mathematically integrated it with the number of colocalizations between the features to define a score for maximal colocalization minimal correlation (MACMIC). The MACMIC score allows us to quantitatively prioritize the feature combinations that have large number of colocalizations but low correlation. We next performed a systematic analysis of MACMIC score between chromatin features using 1,522 datasets for histone modifications or the binding of chromatin proteins from embryonic stem cells as well as somatic cell types. Our analysis successfully recaptured the previously discovered bivalent domain in embryonic stem cells, and further revealed a key role for CCCTC-Binding Factor (CTCF) in the epigenetic regulation of cell identity genes.

## MATERIAL AND METHODS

### Data collection

The RNA-seq, transcript factors and histone modifications ChIP-seq data for human primary cells, human embryonic stem cells and mouse embryonic stem cells were downloaded from GEO database and ENCODE project website (https://www.encodeproject.org/) ^17^. Processed annotated topologically associating domains and loops from HUVEC^40^ were downloaded from GEO. Detailed information of datasets reanalyzed in this study was listed in Table S1 and Table S2.

### Data processing and analysis

Human reference genome sequence version hg19, mouse reference genome sequence version mm9 and UCSC Known Genes were downloaded from the UCSC Genome Browser website ^18^. TPMs of RNA-seq from ENCODE were directly downloaded from ENCODE project. For GEO datasets, RNA-seq raw reads were mapped to the human genome version hg19 using TopHat version 2.1.1 with default parameter values. The expression value for each gene was determined by the function Cuffdiff in Cufflinks version 2.2.1 with default parameter values.

For ChIP-seq data, reads were first mapped to reference genome by Bowtie version 1.1.0. Peak calling and generation of .wig file were performed by DANPOS 2.2.3. Bigwig was generated using the tool WigToBigWig. The tool WigToBigWig was downloaded from the ENCODE project website (https://www.encodeproject.org/software/wigtobigwig/) ^17^. Then bigwig file was submitted to the UCSC Genome Browser (https://genome.ucsc.edu) to visualize the ChIP-Seq signal at each base pair ^18,19^. The average density plots of epigenetic marks in promoter region around TSS were plotted using the Profile function in DANPOS version 2.2.3. Heatmap was plotted using Morpheus (https://software.broadinstitute.org/morpheus). P values of boxplots were calculated with a two-sided Wilcoxon test. For the regulation network, we used CellNet method^20^ to define the network and downloaded the network nodes (genes), edges and value of closeness between nodes from CellNet website (http://cellnet.hms.harvard.edu/). As the gene number will affect the percentage and p value of overlap between gene groups, we used the same number of top genes from each group to avoid this effect. Because the genes associated with Broad H3K4me3 was reported to be around 500 in each cell type ^21^, we used this number of genes for each gene group.

### Integrated analysis of two chromatin features

For individual markers, the ranking of genes was based on the width of individual markers on the gene promoter region (upstream 3kb of TSS to downstream 10kb of TSS). For the ranking of genes based on the colocalization of two chromatin features, the rank product of two individual markers was calculated first. We defined rank product as 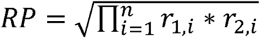, where the *r*_1,*i*_, is the rank of wide for the first marker, the *r*_2,*i*_,, is the rank of wide for the second marker. Then if no colocalization of these two chromatin markers was detected in the gene promoter region, the gene was being removed from the ranking. A colocalization of two chromatin markers at a specific genomic locus was defined by requiring at least 1bp overlap. To measure the colocalization level of two chromatin markers, we calculated the total number of genomic loci that display overlap of these two chromatin markers across whole genome. Afterward, the genes associated with the colocalization of these two chromatin features were ranked based on the rank product of individual features. For a fair comparison, each group defined by H3K4me3, H3K27ac, H3K27me3, colocalization of broad H3K4me3 and broad H3K27me3, colocalization of broad H3K4me3 and broad H3K27ac contained only the top 500 genes. GO term pathway analysis was performed by the web portal (http://geneontology.org/)^22^.

### Calculation of MACMIC score

To calculate MACMIC score, we first calculated Mutual information (MI) that is a widely used measure of the mutual dependence between two variables. More specifically, MI measures how much does the knowledge of one variable reduces uncertainty of the other variable. If two chromatin markers have larger MI, these two chromatin markers share more information and are less independent from each other. Mathematically, mutual information is calculated by following equations:

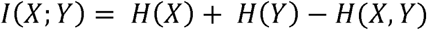

where *X* and *Y* represent the peak width from two different chromatin features, *I(X;Y)* is the mutual information of *X* and *Y. H(X)* and *H(Y) are* the marginal entropies and *H(X,Y)* is the joint entropy of *X* and *Y.* Entropies are calculated by the following equation:

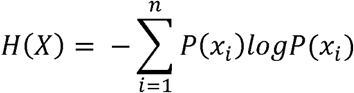

where *n* is the total gene number, *P(x*_*i*_) is the probability by which the total signal of a given genomic marker is *x*_*i*_ in the promoter region of gene *i*. To calculate *H(X)*, we focused on the promoter region from 3kb upstream to 10kb downstream of transcription start site. For a promoter that has multiple ChiP-seq peaks, we calculated the total signal that is the sum of signals in these peaks. The function Selector in DANPOS was used to map peaks to promoters. And we used Poisson distribution to calculate the probability of the observed ChiP-seq signal in a given promoter region ^23^. To calculate the joint entropy of two genomic features, we used the following equation:

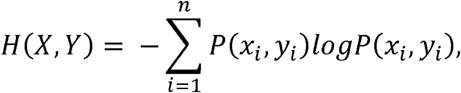

where *n* is the total gene number, *P(x*_*i*_, *y*_*i*_) is the joint probability that the total signals of the first and second markers are *x*_*i*_ and *y*_*i*_ in the promoter region of gene *i*.

Considering the penalty of high correlation feature pairs, MACMIC score is calculated by the following equation:

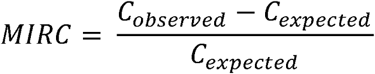

where C represents the colocalization of two chromatin features which is counted by the number of overlap events. The p-value for each term tests the null hypothesis that the residual is equal to zero. A low p-value (<0.05) indicates that for a specific value of MI, the feature combinations have a significant higher colocalization than the estimated colocalization on the genome.

### CTCF associated super-enhancers

CTCF ChIP-seq datasets were processed as previously described. Peaks with height larger than upper quartile of peak height values were defined as high confidence CTCF peaks. Super-enhancers were defined as previous defined^24^, and then super-enhancers were categorized into two categories based on the existence of high confidence CTCF peaks within super-enhancers. Super-enhancers with high confidence CTCF peaks were named as CTCF associated super-enhancers (CSEs). Super-enhancers without high confidence CTCF peaks were named as CTCF associated super-enhancers (OSEs).

### Simulation of association between CTCF and enhancers

For each group of typical enhancers, each typical enhancer was randomly matched to a super-enhancer and then typical enhancers were enlarged towards two directions until they had the same size as super-enhancers. Associations of CTCF and super-enhancers, typical enhancers and enlarged typical enhancers were calculated based on the overlap events between these two different epigenetic markers.

## RESULTS

### Colocalization of globally low-correlation chromatin features reveals unique functional pathways

We first tested whether the colocalization of two histone modifications could identify genes that were not effectively identified by each of the two modifications. We performed the analysis for H3K4me3 and H3K27ac that had strong correlation across the genome (**Fig. S1A**) and compared it to the analysis for H3K4me3 and H3K27me3 that had little correlation across the genome (**Fig. S1B**) in human stem cell H1. We recently revealed that the top 500 genes associated with broad H3K4me3 were enriched with tumor suppressor genes^21^. For a fair comparison, we retrieved the top 500 genes associated with broad H3K27ac and the top 500 genes associated with broad H3K27me3. There were 288 (57.6%) genes associated with both broad H3K4me3 and broad H3K27ac (**Fig. S1C**). In contrast, there was no gene associated with both broad H3K4me3 and broad H3K27me3 (**Fig. S1D**). To further explore the potential colocalization between H3K4me3 and H3K27me3, we defined the top 500 genes by the rank product of H3K4me3 width and H3K27me3 width (H3K4me3&H3K27me3 broad colocalization) (**Fig. S1E**). We also defined the top 500 genes by the rank product of H3K4me3 width and H3K27ac width (H3K4me3&H3K27ac broad colocalization) (**Fig. S1E**). For the genes associated with H3K4me3&H3K27ac broad colocalization, only 7 genes were not captured by broad H3K4me3 or broad H3K27ac (**Fig. S1C**). However, for the genes associated with H3K4me3&H3K27me3 broad colocalization, 421 (84.2%) genes were not captured by broad H3K4me3 or broad H3K27me3 (**Fig. S1D**). Further, for the 2168 pathways significantly enriched in genes associated with H3K4me3&H3K27me3 broad colocalization, 1404 pathways showed no significant enrichment in genes associated with broad H3K4me3 or broad H3K27me3 (**Fig. S1F**). These pathways were mainly related to somatic cell lineage specification (**Fig. S1G**), which agreed with the reported role of bivalent domains. These results suggested that colocalization of globally low-correlation features in a cell could be associated with unique biological implications that were not associated with the localization of each of these features.

### A new method to identify features with maximal colocalization minimal correlation (MACMIC)

Here, we used mutual information as an indication for correlation because mutual information is more general than other methods such as linear correlation. A large mutual information value will indicate strong correlation that can be either positive or negative, and either linear or nonlinear. Theoretically, two features that have a small mutual information value tend to have no or a small number of colocalization, whereas a large number of colocalizations are often associated with large mutual information value. To develop a simple method to prioritize feature pairs that have minimal correlation but a maximal number of colocalizations, we first performed a systematic analysis of the relationship between the mutual information value and the number of colocalizations for 225 feature pairs derived from 6 chromatin features in 15 cell types (**Table S1**). We analyzed 6 features, which formed 15 pairs with each other in each cell type and thus resulted in 225 feature pairs in 15 cell types (**Table S3**). Most feature pairs displayed a positive correlation between the mutual information value and the number of colocalizations (Spearman correlation coefficient 0.46) (**Fig. 1A**). Similar results were observed by replacing mutual information with absolute value of correlation coefficient or PCA value (**Fig. S2**). However, there were a few feature pairs that displayed a large number of colocalizations but small mutual information value (**Fig. 1A**). We therefore developed a regression model that employed the mutual information value to determine an expected number of colocalizations, and next utilized the normalized discrepancy between the observed and the expected numbers of colocalizations as a measurement of the MACMIC (**Fig. 1B**). We calculated the MACMIC scores for the 225 individual feature pairs and found that the large MACMIC scores effectively prioritized feature pairs that possessed large number of colocalizations but weak correlations (**Fig. 1C**). We further tested our MACMIC analysis method on 3160 feature pairs derived from 80 chromatin features in human ESC H1. Our result again indicated that MACMIC successfully prioritized the feature pairs with minimal mutual information but substantial colocalizations **(Fig. 1D)**.

**Figure 1.**
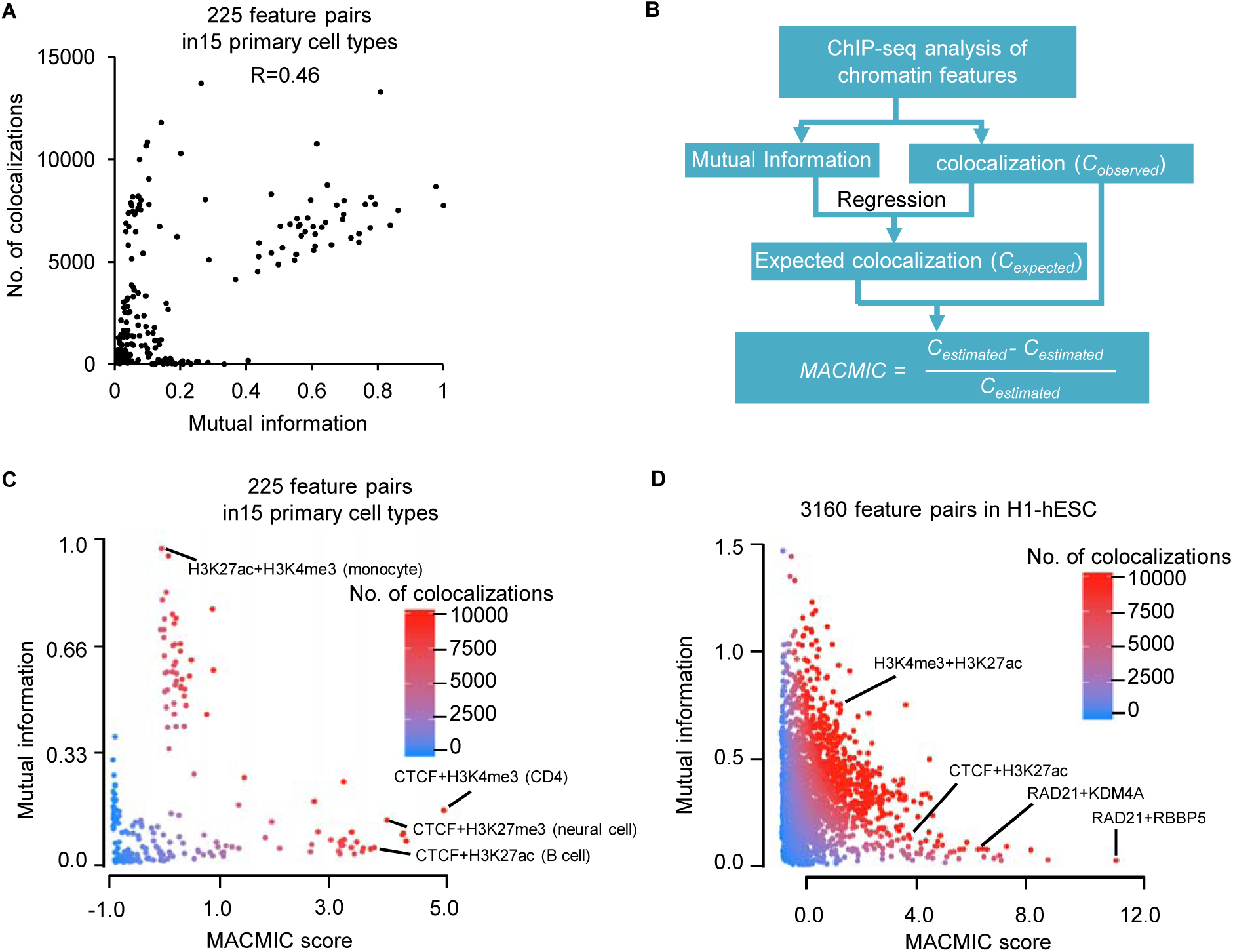
The MACMIC method to define mutual information redundancy of colocalizations between genomic features. (**A**) Scatter plot to show mutual information value and the number of colocalization for each of 225 feature pairs derived from 6 features that form 15 combinations with each other in each of 15 human primary cell types. **(B)** The workflow to calculate the MACMIC score. (**C**) Scatter plot to show MACMIC score and mutual information value for each of 225 feature pairs derived from 6 features that form 15 combinations with each other in each of 15 human primary cell types. Color scale indicates the number of colocalizations between each pair of features. (**D**) Scatter plot to show MACMIC score and mutual information value between each pair of features. 3160 feature pairs derived from 80 features in H1-hESC were plotted. Color scale indicates the number of colocalizations between each pair of features.

### MACMIC identifies a unique association of CTCF with super-enhancer

To further test whether MACMIC scores could effectively recapture feature pairs with biological implications, we analyzed MACMIC scores between H3K4me3 and H3K27me3 in 15 human primary cell types. In agreement with the reported large number of bivalent domains marked by both H3K4me3 and H3K27me3 in embryonic stem cells, we observed a large MACMIC score (2.8) in the H1 cell. On the other hand, in agreement with the reported resolution of bivalent domains to form either repressive domains marked by H3K27me3 or active domains marked by H3K4me3, the MACMIC scores between H3K4me3 and H3K27me3 were low in all the 14 somatic cell types (from −0.68 to 0.67) **(Fig. 2A).** Therefore, MACMIC analysis successfully recaptured bivalent domains that were known to play a key role in embryonic stem cells.

**Figure 2.**
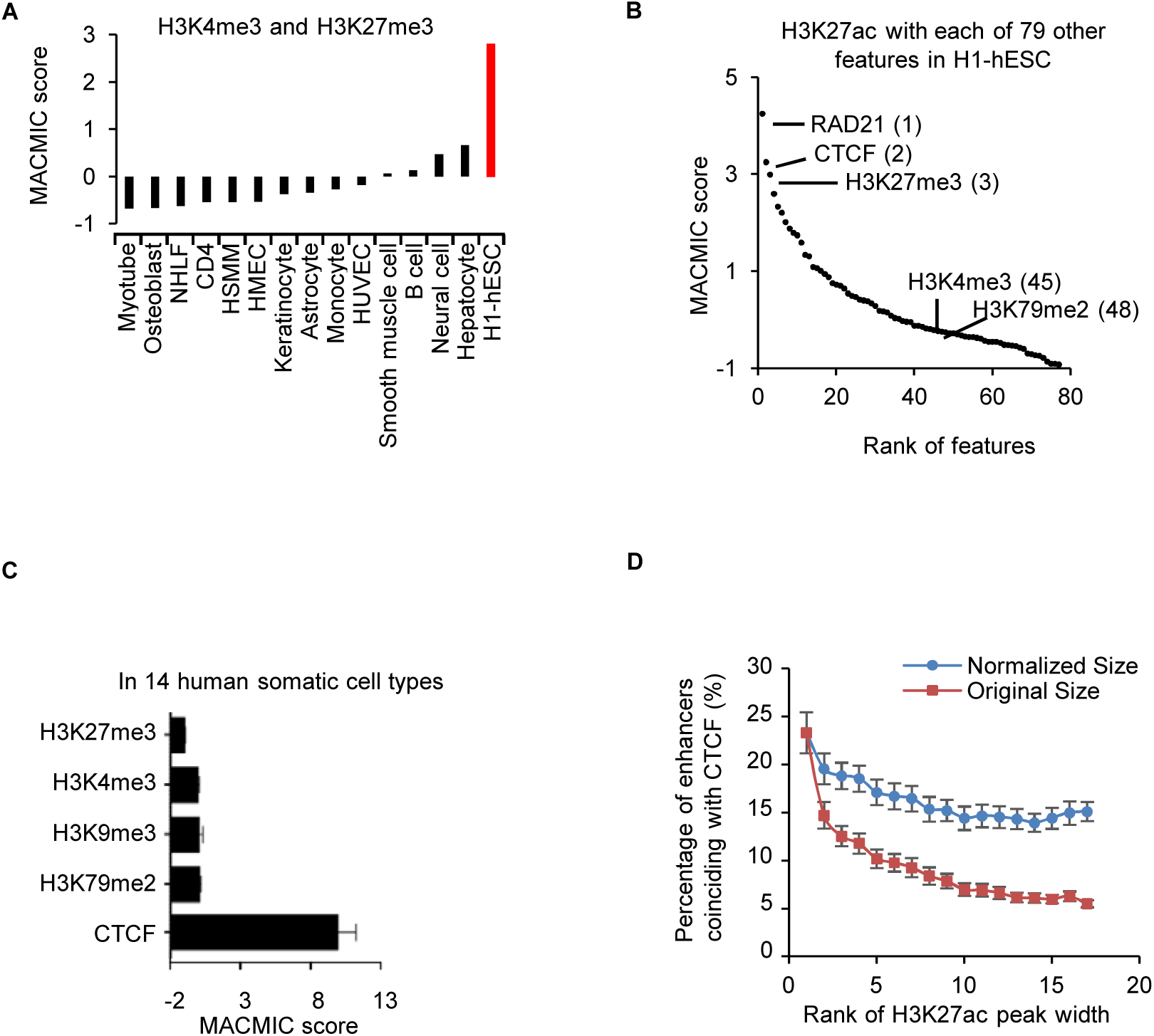
MACMIC reveals minimal information redundancy of frequent colocalizations between CTCF binding sites and super-enhancers. **(A)** Bar plot to show MACMIC scores between H3K4me3 and H3K27me3 in individual human primary cell types. (**B**) MACMIC scores between H3K27ac and individual other chromatin features in H1-hESC. (**C**) MACMIC scores between H3K27ac and individual other chromatin features in 14 human somatic primary cell types. Error bars indicate the standard deviation of MACMIC scores across cell types. (**D**) Percentage of enhancers that coincided with CTCF binding sites in 15 human primary cell types. Enhancers were divided into individual groups on the base of their H3K27ac width. Each group contains 500 enhancers, e.g. rank 1 contains the widest 500 enhancers; rank 2 contains the 501st to 1000th widest enhancers.

We next tested whether MACMIC analysis could successfully identify new feature pairs that possess a large number of functionally important colocalizations but low correlation. We ranked a set of 79 chromatin features in H1 cells by the MACMIC scores between the enhancer feature H3K27ac and each of these features **(Fig. 2B**). The top features with the large MACMIC scores in the rank included the suppressive histone modification H3K27me3, consistent with the implication that H3K27ac and H3K27me3 might co-exist in bivalent domains. Interestingly, master regulators of three-dimensional chromatin interaction, the CTCF^25^ and its binding partner RAD21^26^, topped in the rank list (**Fig. 2B**). We further performed analysis in 14 human somatic cell types that each had ChIP-seq datasets for a set of 6 chromatin features from the ENCODE project^17^ (**Table S1**). The results showed that the MACMIC score between H3K27ac and the binding of CTCF was significantly larger than MACMIC scores between H3K27ac and the other 4 features including H3K27me3, H3K4me3, H3K9me3 and H3K79me2 (**Fig. 2C**). Moreover, colocalization analysis for CTCF and H3K27ac found that CTCF binding sites had the largest number of colocalizations with the broadest H3K27ac peaks (super-enhancers) (**Fig. 2D**). To test whether this higher frequency of colocalization was simply due to the longer DNA sequences of super-enhancers, we performed a normalization by lengthening typical enhancers at the two ends of each enhancer, so that the DNA sequences assigned to typical enhancers had equivalent sizes to those of super-enhancers. The result showed that the frequency of colocalization with CTCF binding sites still tended to be higher for super-enhancers when compared to other enhancers (**Fig. 2D**).

### A unique enrichment of CTCF associated super-enhancer in cell identity genes

Since super-enhancers were reported to regulate cell identity genes^24^, we determined to investigate the role of CTCF in this regulation. We divided super-enhancers into two categories, i.e., CTCF associated super-enhancers (CSEs) and other super-enhancers (OSEs). To study the function of genes marked by CSEs and OSEs, we defined the genes of which the gene body overlapped with CSEs (or OSEs) for at least 1bp as the CSEs (or OSEs) marked genes Intriguingly, only the genes marked by CSEs were significantly enriched in the pathways associated with cell lineage specifications, e.g., the endothelial cell differentiation pathway (GO:0045601) for CSEs in human umbilical vein endothelial cells (HUVECs) (**Fig. 3A**) and the neuron differentiation pathway (GO:0045664) for CSEs in neural cells (**Fig. 3B**). Manual inspection of individual known cell lineage factors in these cell types further confirmed the colocalization of ChIP-seq signals of H3K27ac and CTCF, e.g., at the gene NR2F2^27^ in endothelial cells and the gene FOXG1^28^ in neural cells (**Fig. 3C, D**). In contrast, some other genes, although also displaying broad enrichment of H3K27ac, were depleted of CTCF binding sites, e.g., at the gene ARF1 in endothelial cells and the gene PON1 in neural cells (**Fig. 3C, D**). Intriguingly, there were typically multiple binding sites of CTCF located within the active region of each CSE. This colocalization pattern was different from the well-known function of CTCF binding sites as insulators, which often happened between active domain and repressive domain. Besides, a significant portion of the genes associated with CSEs encoded transcription factors, whereas we did not observe this phenomenon for the genes associated with OSEs (**Fig. 3E**). Further, the genes associated with CSEs were connected to a significantly large number of edges in the gene regulatory networks, whereas the numbers of connected network edges were similar for genes associated with OSEs and random control genes (**Fig. 3F**). The differences between CSEs and OSEs in their association with genes in cell lineage pathways were highly reproducible in the other 15 primary cell types that we have analyzed (**Fig. 3G**). It was reported that the establishment of cell type specific chromatin loops were important during cell differentiation^29^. Consistently, we found that CSEs were enriched near chromatin loops (**Fig. S3A**) and the boundaries of topologically associating domains (TADs) (**Fig. S3B**), whereas no significant differences in the sizes of the associated TADs were observed between CSEs and OSEs (**Fig. S3C**).

**Figure 3.**
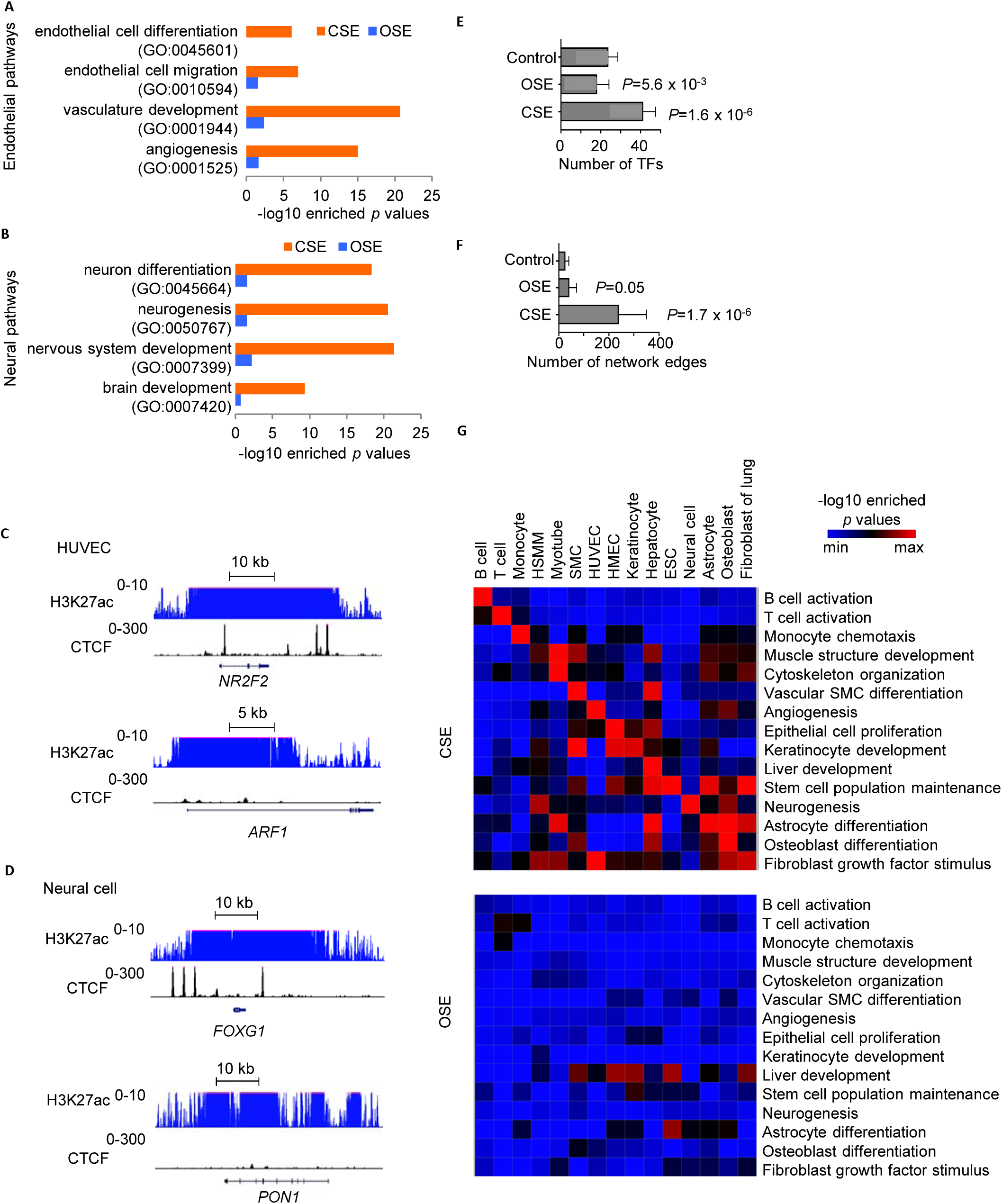
CTCF-associated super-enhancers mark cell identity genes. (**A-B**) Individual pathways enriched in CSEs or OSEs genes in HUVECs **(A)** or neural cells **(B)**. (**C-D**) ChIP-Seq signals for H3K27ac and CTCF at CSE gene NR2F2 and OSE gene ARF1 in HUVECs (**C**) and CSE gene FOXG1 and OSE gene PON1 in neural cells (**D**). (**E-F**) The number of transcription factors within each gene group **(E)** and the number of network edges within each gene group **(F)** in 15 human primary cell types. Error bars indicate the standard deviation across cell types. Each gene group was defined to have the same number of genes. P values were determined by Wilcoxon test in comparison to the control group. (**G**) Heatmap to show −log10 enrichment P values of cell type related pathways (rows) in CSEs genes (top panel) or OSEs genes (bottom panel) defined in each cell type (columns).

### CSE- and OSE-associated genes have similar expression levels and cell type specificities

To understand how CTCF regulates enhancer activity and in turn regulates cell identity, we first compared the expression levels of associated genes between CSEs and OSEs. Intriguingly, similar expression levels were observed between CSE-marked genes and OSE-marked genes, and this result was highly reproducible in HUVECs (**Fig. 4A left panel**) and neural cells (**Fig. 4A right panel**). Further, CSEs genes and OSEs genes of HUVECs were both significantly up regulated in HUVECs compared to embryonic stem cells and neural cells (**Fig. 4B left panels**). Consistently, CSEs genes and OSEs genes of neural cells were both significantly up regulated in neural cells compared to embryonic stem cells and HUVECs (**Fig. 4B right panels**). These results suggested that CSEs and OSEs genes of the same cell type had similar expression levels and cell type specificities.

**Figure 4.**
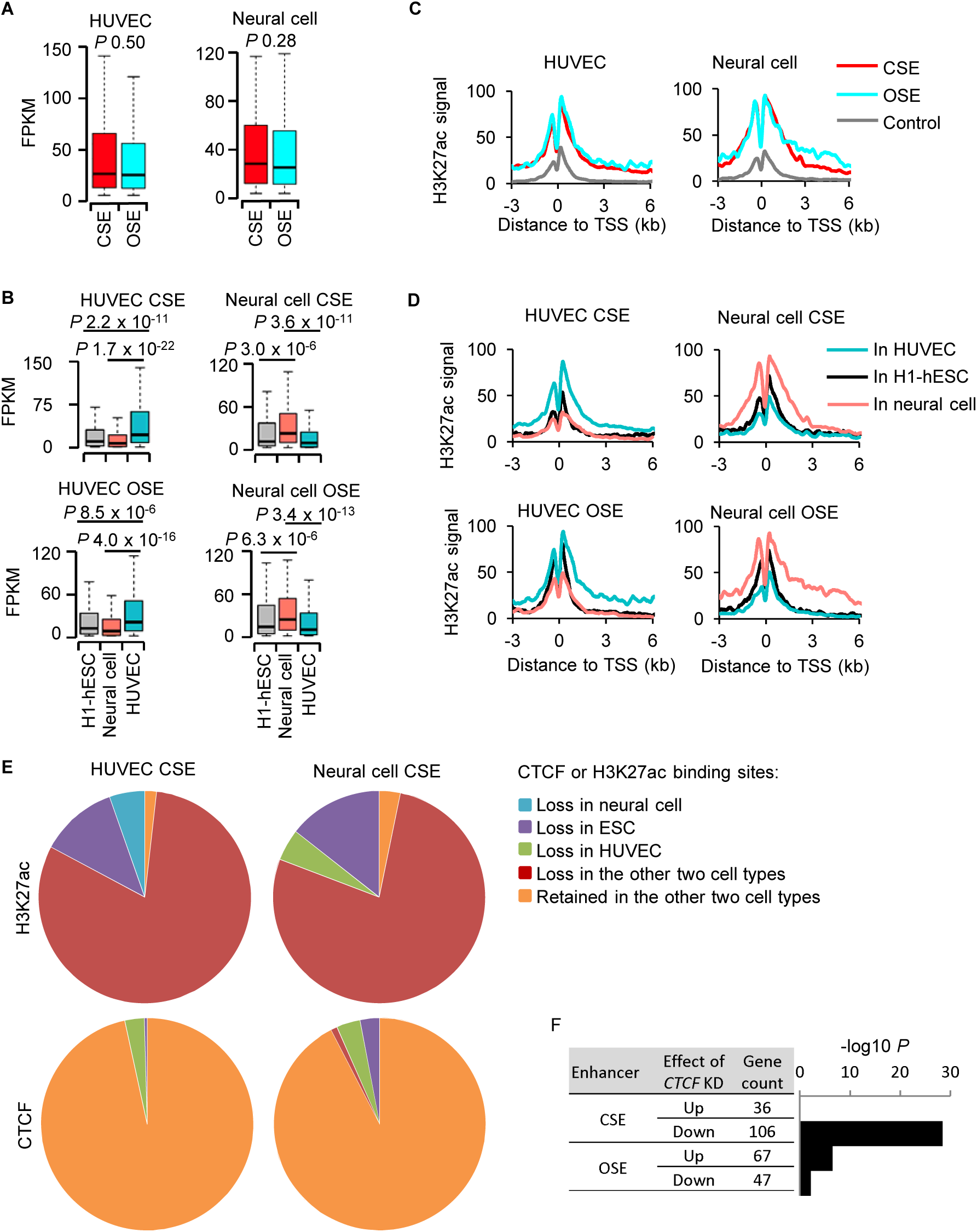
CTCF is linked to the activation of enhancers. (**A**) Box plot to show RNA expressions of HUVEC (left panel) and neural cell (right panel) CSEs genes and OSEs genes in cell types that defined them. (**B**) Box plot to show RNA expressions of CSE genes (top panels) and OSE genes (bottom panels) in neural cell, HUVEC, and H1-hESC. CSEs and OSEs were defined in HUVEC (left panels) or neural cells (right panels) (**C**) H3K27ac signals at HUVEC (left panel) and neural cell (right panel) CSEs genes, OSEs genes, and control genes in the cell type that defined these gene groups. **(D)** H3K27ac signals at CSE genes (top panels) and OSE genes (bottom panels) in HUVEC, H1-hESC, and neural cells. CSEs and OSEs were defined in HUVEC (left panels) or neural cells (right panels). **(E)** Pie charts to show binding status of CTCF at HUVEC CSEs in neural cells or embryonic stem cells (top left), binding status of CTCF at neural cell CSEs in HUVEC or embryonic stem cells (top right), H3K27ac status at HUVEC CSEs in neural cells or embryonic stem cells (bottom left), and H3K27ac status at neural cell CSEs in HUVEC or embryonic stem cells (bottom right). (**F**) Barplot to show –log10 enrichment P values of CSE genes or OSE genes in the genes up or down regulated by shCTCF in HeLa cells. P values were determined by Wilcoxon test (A, B, E, F).

We next compared the H3K27ac levels between CSEs and OSEs, as H3K27ac is a marker for enhancer activation. The result indicated that the H3K27ac levels were similar at CSEs and OSEs within HUVECs (**Fig. 4C left**). Similarly, the H3K27ac levels were similar at CSEs and OSEs within neural cells (**Fig. 4C right**). Further, the H3K27ac levels at HUVEC-specific CSEs and OSEs were higher in HUVECs when compared to the same regions in embryonic stem cells and neural cells, whereas the H3K27ac levels at neuron-specific CSEs and OSEs were higher in neural cells compared to the same regions in HUVECs and embryonic stem cells (**Fig. 4D**). Therefore, in agreement with result from the expression analysis, CSEs and OSEs genes of the same cell type had similar epigenetic states and specificities.

Of the top 500 HUVEC CSEs, 405 (81%) lost H3K27ac in neural cells and embryonic stem cells (**Fig. 4E top left)**. In contract, the binding of CTCF in 483 (97%) HUVEC CSEs were retained in both neural cells and embryonic stem cells (**Fig. 4E bottom left**). Similar results were observed for the neural cell CSEs. Of the top 500 neural CSEs, 388 (78%) lost H3K27ac in HUVECs and embryonic stem cells (**Fig. 4E top right)**, while the binding of CTCF in 462 (92%) neural cell CSEs were retained in both HUVECs and embryonic stem cells (**Fig. 4E bottom right**). To further understand the role of CTCF in CSEs, we next analyzed an RNA-Seq dataset from HeLa cells with CTCF knocked down or not. The genes associated with CSEs of HeLa cells were significantly enriched in the genes down regulated but not in the genes up regulated in response to CTCF knockdown (**Fig. 4F**). In contrast, the OSEs associated genes showed little enrichment in the down or up regulated genes induced by knockdown of CTCF (**Fig. 4F**). Of the top 500 HUVEC OSEs, 331 (66%) lost H3K27ac in neural cells and embryonic stem cells (**Fig. S4 top left)**. In contract, the binding of CTCF in 492 (98%) HUVEC OSEs were retained in both neural cells and embryonic stem cells (**Fig. S4 bottom left**). Similar results were observed for the neural cell OSEs. Of the top 500 neural OSEs, 347 (69%) lost H3K27ac in HUVECs and embryonic stem cells (**Fig. S4 top right)**, while the binding of CTCF in 476 (96%) neural cell OSEs were retained in both HUVECs and embryonic stem cells (**Fig. S4 bottom right**). These results indicated that although the loss of the activation state of CSEs may not require the loss of CTCF bindings, the bindings of CTCF were required for the activation of CSEs and their associated genes.

### CSEs of a given cell type display increased repressive modification H3K27me3 in other cell types

A cell identity gene has two key attributes: 1) it is associated with active chromatin modifications and thus activated to play an important role in the cell type that requires its activation; and 2) it is associated repressive chromatin modifications and thus silenced in most other cell types. Since our results demonstrated that the CSEs of one cell type lost H3K27ac but retained the binding of CTCF in other cell types, we hypothesized that the binding of CTCF might be also important for the repressions of these CSEs in the other cell types.

We first defined CSEs, OSEs, and a set of random control genes in HUVECs, and analyzed the pattern of the repressive histone modification H3K27me3 on these 3 gene sets in each of three cell types including embryonic stem cells, neural cells and also HUVECs. We found that the H3K27me3 signals in HUVEC showed a similar pattern at the HUVEC CSEs as at the HUVEC OSEs, but are substantially weaker than at the random control genes (**Fig. 5A top**). Intriguingly, only the CSEs of HUVECs, not the OSEs of HUVECs or the random control genes, were marked by strong H3K27me3 signals in embryonic stem cells (**Fig. 5A middle**). These trends observed for H3K27me3 in embryonic stem cells were the same for H3K27me3 in neural cells (**Fig. 5A bottom**). Similar results were observed when we defined CSEs, OSEs, and a set of random control genes in neural cells to analyze the pattern of H3K27me3 on these 3 gene sets in HUVECs, embryonic stem cells, and neural cells. The H3K27me3 signals in neural cells showed a similar pattern at the neural CSEs as at the neural OSEs, but are substantially weaker at the random control genes (**Fig. 5B bottom**). However, only the CSEs of neural cells, not the OSEs of neural cells or the random control genes, possessed strong H3K27me3 signals in embryonic stem cells (**Fig. 5B middle**). These trends observed for H3K27me3 in embryonic stem cells were the same for H3K27me3 in HUVECs (**Fig. 5B top**).

**Figure 5.**
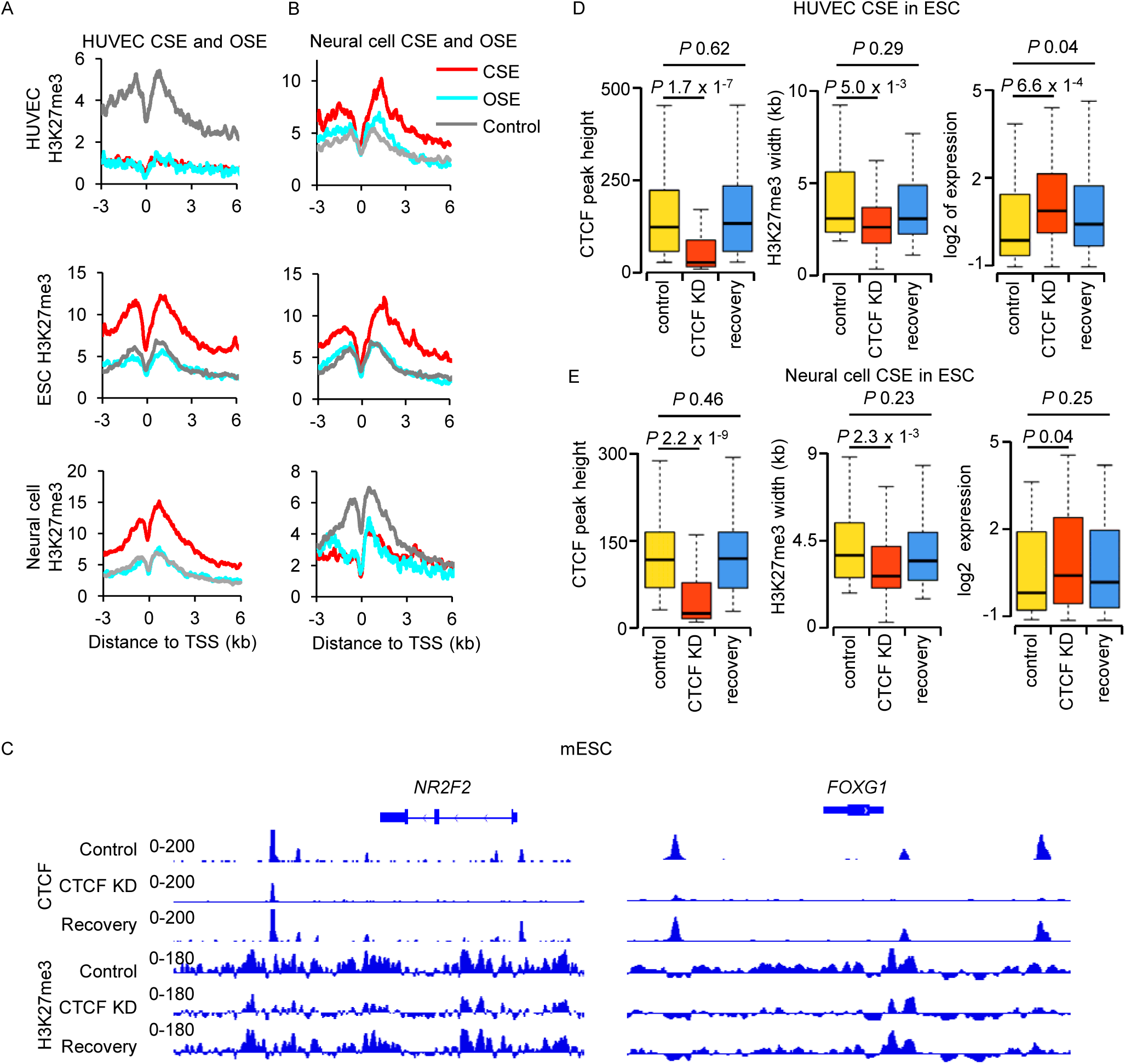
CTCF regulates cell identity by facilitating the suppressive marker H3K27me3. (**A-B**) H3K27me3 signals in H1-hESC, neural cell and HUVECs at CSE genes, OSE genes and control genes defined in HUVECs (**A**) and neural cells **(B). (C)** ChIP-Seq signals for CTCF and H3K27me3 in mESC at the HUVEC CSE gene *NR2F2* (left) and the neural CSE gene *FOXG1* (right). **(D-E)** Box plot to show the heights of CTCF ChIP-Seq enrichment peaks, the widths of H3K27me3 enrichment domains, and the RNA expressions of CSE genes of HUVECs (**D**) and CSE genes of neural cells **(E)** under different conditions in embryonic stem cells. P values were determined by Wilcoxon test.

We next further expanded our analysis to 15 sets of biosamples that each had ChIP-Seq data for CTCF, H3K27ac, and H3K27me3. Consistent with the results from HUVECs and neural cells, CSEs and OSEs showed similar enrichment of H3K27ac (**Fig. S5A**) and similar depletion of H3K27me3 (**Fig. S5B**) in cell types that defined these CSEs and OSEs. Next, we analyzed these CSEs and OSEs in H3K27ac ChIP-Seq datasets from 84 biosamples and H3K27me3 ChIP-Seq datasets from 125 biosamples from the ENCODE database. CSEs and OSEs both showed attenuated enrichment of H3K27ac when the H3K27ac was analyzed in cell types different from the cell types that defined the CSEs and OSEs (**Fig. S5C**). However, the CSEs were associated with significant enrichment of H3K27me3, whereas the OSEs showed little enrichment of H3K27me3, when the H3K27me3 was analyzed in cell types different from the cell types that defined these CSEs and OSEs (**Fig. S5D**). These analyses indicated that the CSEs, but not the OSEs, were under stringent epigenetic repression by H3K27me3 in cell types different from the cell types that defined the CSEs and OSEs. Interestingly, CTCF and H3K27me3 is also among the top feature pairs ranked by MACMIC score in H1 hESC (**Fig. S6**).

### CTCF in a given cell type is required for the repression of genes associated with the CSEs defined in other cell types

Importantly, auxin-induced degradation of CTCF in embryonic stem cells led to the loss of CTCF bindings and H3K27me3 signals in embryonic stem cells at the CSEs genes defined in HUVECs as well as the CSEs genes defined in neural cells. For example, signals of CTCF bindings and H3K27me3 in embryonic stem cells at known identity genes of somatic cell types, the NR2F2^27^ of endothelial cells (**Fig. 5C left**) and the FOXG1^28^ of neural cells (**Fig. 5C right**), were substantially attenuated after auxin induced degradation of CTCF, and recovered after auxin was washed off (**Fig. 5C**). The CTCF binding sites in embryonic stem cells at these genes were located within the broad H3K27me3 modifications. This colocalization of CTCF binding sites and broad H3K27me3 in embryonic stem cells was similar to the colocalization observed for CTCF binding sites and super-enhancers in HUVECs, neural cells, heart, fibroblast cell, and bone marrow macrophage cell. Our further analysis indicated that in parallel with the loss of CTCF bindings in embryonic stem cells at these genes (**Fig. 5D left and Fig. 5E left**), the H3K27me3 signals in embryonic stem cells were reduced dramatically (**Fig. 5D, Fig. 5E, Fig. S7 middle**) and the expressions in embryonic stem cells were significantly up regulated (**Fig. 5D, Fig. 5E, Fig. S7 right**). Taken together, these results suggested that the CTCF in a given cell type was required for the repression of genes associated with the CSEs defined in a different cell type.

## DISCUSSION

Conventional analysis of relationship between chromatin features tends to focus on strongly positive or negative correlation to identify the associated components within a specific biological process^1^. However, genomic features with weak correlation across the genome may still colocalize at many genomic sites in a biologically important manner. It was hard to capture the significance of such colocalizations on the basis of conventional correlation analysis. In this study, we provide a new method to identify MACMIC, which effectively prioritize the feature pairs with low genome-wide correlation but substantial colocalizations. Using the MACMIC, we successfully recapture the reported bivalent domain in embryonic stem cells, which is composed of both activating histone modifications, e.g., H3K4me3, and the repressive histone modifications, e.g., H3K27me3. Activating histone modification and the repressive histone modification possess low genome-wide correlation in the embryonic stem cell, but the colocalizations of them at bivalent domains mark important lineage specific regulators.

As proof of principle, we present a novel relationship identified by MACMIC between the bindings of CTCF and the enhancer marker H3K27ac. Our analysis demonstrated that their colocalization is key to both the activation and repression of cell identity genes. Numerous efforts have been made to understand cell identity regulation^20^. Somatic cells such as fibroblasts^30^, keratinocytes^31^, peripheral blood cells^32^, and neural progenitor cells^33^, have been sucessfully reprogrammed to induced pluripotent stem cells. Many transcription factors and epigenetic regulators have been proposed to play important roles in these dynamic processes. We and several other groups recently discovered that cell identity genes manifested unique chromatin epigenetic signatures associated with their distinct transcriptional regulation mechanisms^24,34-36^. CTCF is well known for its function as an insulator that binds between active and repressive domains on chromatin^37^, as a mediator for promoter-enhancer interaction^38^, and as a partner of cohesin in regulating chromatin 3D structure^39,40^. It further has been proven to be an essential factor for cell differentiation and development of T cell^41^, Neuron^42^, Heart^43^, and Limb^44^. However, how these functions of CTCF are connected to the regulation of cell identity genes was not known.

In this study, we separate CSEs from OSEs based on the colocalization of CTCF binding sites with H3K27ac signals in CSEs. Our results suggest that CTCF contributes to the activations of CSEs in cell types that require the activations, and is involved in the repression of CSEs in other cell types that require the repressions. Interestingly, only CSEs showed significantly higher H3K27me3 signals in the cell types that required their repressions, consistent with the notion that epigenetic repression of cell identity genes of a given cell type is critical in other somatic cell types (**Fig. 5, S3**). In response to the loss of CTCF function in embryonic stem cells, H3K27me3 signals in embryonic stem cells at the CSEs of somatic cell types were dramatically reduced but restored after recovery of CTCF function (**Fig. 5**). Intriguingly, the CTCF binding sites in embryonic stem cells at somatic cell identity genes were located within the repressive domains of embryonic stem cells. This colocalization was similar to the colocalization of CTCF binding sites with super-enhancers observed in somatic cell types. These unique CTCF associated epigenetic profiles suggested a novel function of CTCF in epigenetic regulation of transcription.

Recently, many epigenetic regulators have been proven to interact with CTCF in different biological processes. For instance, BRD2 has been reported to directly interact with CTCF during Th17 cell differentiation^45^. This report suggested that CTCF might be able to regulate enhancer signals by facilitating the binding of enhancer mediator on the chromatin^46^. Interestingly, our result indicated that CTCF played an important role for the repressive histone modification, H3K27me3. This observation is consistent with recent reports that depletion of CTCF does not affect the spreading of H3K27me3^47^, indicating that CTCF might affect H3K27me3 modification by a process other than the spreading. Considering that CTCF was reported to regulate Igf2 expression by direct interaction with Suz12, an important component of Polycomb complexes PRC2^48^, it is possible that CTCF may serve as a landmark to facilitate the localization of epigenetic regulators. Due to limited availability of public datasets for human, we defined genes associated with CSEs and OSEs in human HUVEC and neural cells, and analyzed homolog genes in mouse ESC (mESC) with CTCF ChIP-seq and H3K27me3 ChIP-seq data under normal and CTCF-AID conditions. To further validate our results, we used ChIP-Seq data for CTCF and H3K27ac in three mouse primary samples including heart, embryonic fibroblast and bone marrow macrophage to define genes associated with CSEs and OSEs, next analyzed CTCF and H3K27me3 at these genes in mouse ESC (mESC), and still got a consistent result.

Interestingly, among the top-ranked feature pairs in H1-hESC, there are many pairs that each was formed by a factor associated with chromatin structure and a factor associated with histone modification for transcription activation or repression. For example, the combination of RBBP5 ^49^ versus RAD21 ^50^ and the combination of KDM4A ^50^ versus RAD21. RBBP5 and KDM4A are important regulators of H3K4me3, and RAD21 is a component of the cohesion complex that regulates chromatin looping. In addition, we further observed additional combinations that each include a factor associated with activation of transcription and a factor associated with repression of transcription, such as CTBP2 ^51^ and H3K27ac. This kind of combination is consistent with the concept of bivalent domain in the stem cells. Last but not the least, we found high-score combinations that each include a factor of cohesion complex and a factor associated with repression of transcription, such as the combination of CTCF and H3K27me3, which we found later is also very important for the cell identity regulation.

Taken together, through MACMIC analysis, we find that CTCF plays an important role in the epigenetic regulation of cell identity. Further analysis suggests that CTCF is important for the regulation of both enhancer signals and repressive signals at the CSEs in a cell-type specific manner. Although our analysis focused on the colocalization of enhancer signal with the other chromatin features, MACMIC analysis has great potential to identify many other novel biologically significant colocalizations between chromatin features that have low global correlation across the genome. With the increased usage of sequencing technologies, more potential feature pairs can be identified. This will provide opportunities in the future to further understand the function of chromatin in transcription, replication, DNA repairs and many other biological processes.

## AVAILABILITY

All public datasets used in this study are listed in Table S1 for ENCODE database and Table S2 for GEO database. The code for MACMIC is available at the website GitHub, https://github.com/bxia888/MACMIC. The CSE- and OSE-associated genes can be downloaded from **Table S4**.

## SUPPLEMENTARY DATA

Figure S1-5.

Table S1. Datasets reanalyzed from ENCODE.

Table S2. Datasets reanalyzed from GEO.

Table S3. Datasets of MACMIC score and p value.

Table S4. Datasets of CSE- and OSE-associated genes.

## AUTHOR CONTRIBUTIONS

K.C. conceived the project, designed the analysis, and interpreted the data. B.X. and G.W., performed the data analysis. K.C. B.X., and G.W. wrote the manuscript with comments from M.Z., J.L., D.Z., Y.L., Y.B., X.W., and L.Z.

## ACKNOWLEDGEMENT

We would like to thank members of Department of Cardiovascular Sciences at Houston Methodist Research Institute for their thoughtful comments on the project. We further appreciate all researchers who generated the public genomic datasets analyzed in this study.

