## Supplemental figure for "MACMIC Reveals Dual Role of CTCF in Epigenetic Regulation of Cell Identity Genes"

### Slide 1
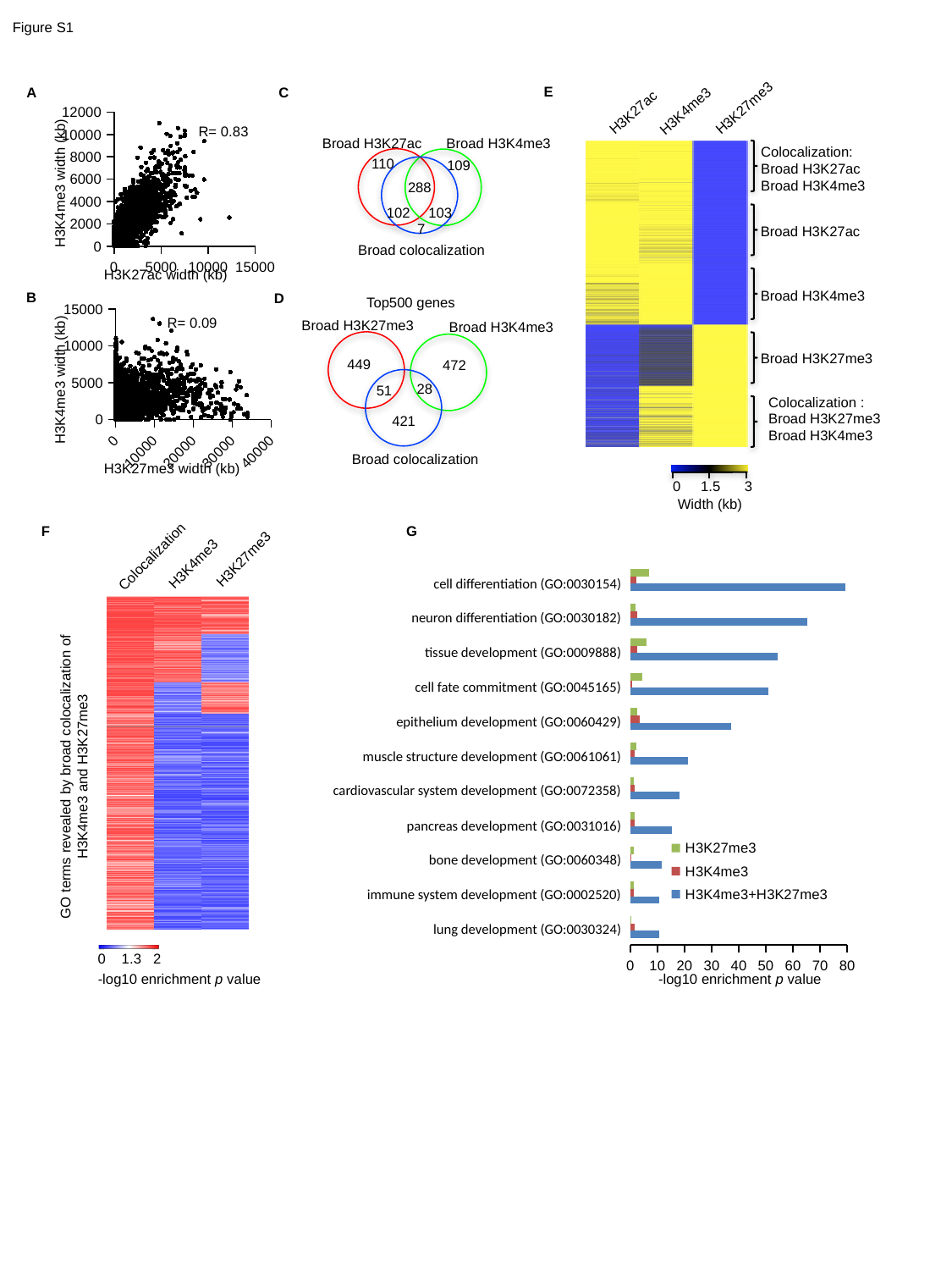

Figure S1
E
C
A
H3K27me3
H3K4me3
H3K27ac
#### Chart
| Category | H3K4me3_total_width |
|---|---|R= 0.83
Broad H3K27ac
Broad H3K4me3
110
109
288
102
103
7
Broad colocalization
Colocalization:
Broad H3K27ac
Broad H3K4me3
H3K4me3 width (kb)
Broad H3K27ac
H3K27ac width (kb)
Broad H3K4me3
B
D
Top500 genes
Broad H3K27me3
Broad H3K4me3
449
472
28
51
421
Broad colocalization
#### Chart
| Category | H3K4me3_total_width |
|---|---|R= 0.09
Broad H3K27me3
H3K4me3 width (kb)
Colocalization :
Broad H3K27me3
Broad H3K4me3
H3K27me3
H3K4me3
Colocalization
GO terms revealed by broad colocalization of H3K4me3 and H3K27me3
0 1.3 2
-log10 enrichment p value
H3K27me3 width (kb)
0 1.5 3
#### Chart
| Category | H3K4me3+H3K27me3 | H3K4me3 | H3K27me3 |
|---|---|---|---|
| lung development (GO:0030324) | 10.5800442515102 | 1.70114692359029 | 0.359518563029578 |
| immune system development (GO:0002520) | 10.6516951369518 | 1.36051351073141 | 1.24949160514865 |
| bone development (GO:0060348) | 11.6556077263148 | 0.329754146925876 | 1.31875876262441 |
| pancreas development (GO:0031016) | 15.3098039199714 | 1.52432881167557 | 1.56703070912559 |
| cardiovascular system development (GO:0072358) | 17.99139982823798 | 1.76447155309245 | 1.30016227413275 |
| muscle structure development (GO:0061061) | 21.2740883677049 | 1.71444269099222 | 2.38933983691012 |
| epithelium development (GO:0060429) | 37.197226274708 | 3.36051351073141 | 2.489454989793379 |
| cell fate commitment (GO:0045165) | 51.0092173081968 | 0.568636235841012 | 4.342944147142886 |
| tissue development (GO:0009888) | 54.34390179798708 | 2.465973893943858 | 5.896196279044036 |
| neuron differentiation (GO:0030182) | 65.28988263488803 | 2.472370099128657 | 1.79588001734407 |
| cell differentiation (GO:0030154) | 79.2588484011482 | 2.27408836770495 | 6.79588001734407 |Width (kb)
F
G
| cell differentiation (GO:0030154) |
| --- |
| neuron differentiation (GO:0030182) |
| tissue development (GO:0009888) |
| cell fate commitment (GO:0045165) |
| epithelium development (GO:0060429) |
| muscle structure development (GO:0061061) |
| cardiovascular system development (GO:0072358) |
| pancreas development (GO:0031016) |
| bone development (GO:0060348) |
| immune system development (GO:0002520) |
| lung development (GO:0030324) |
-log10 enrichment p value

### Slide 2
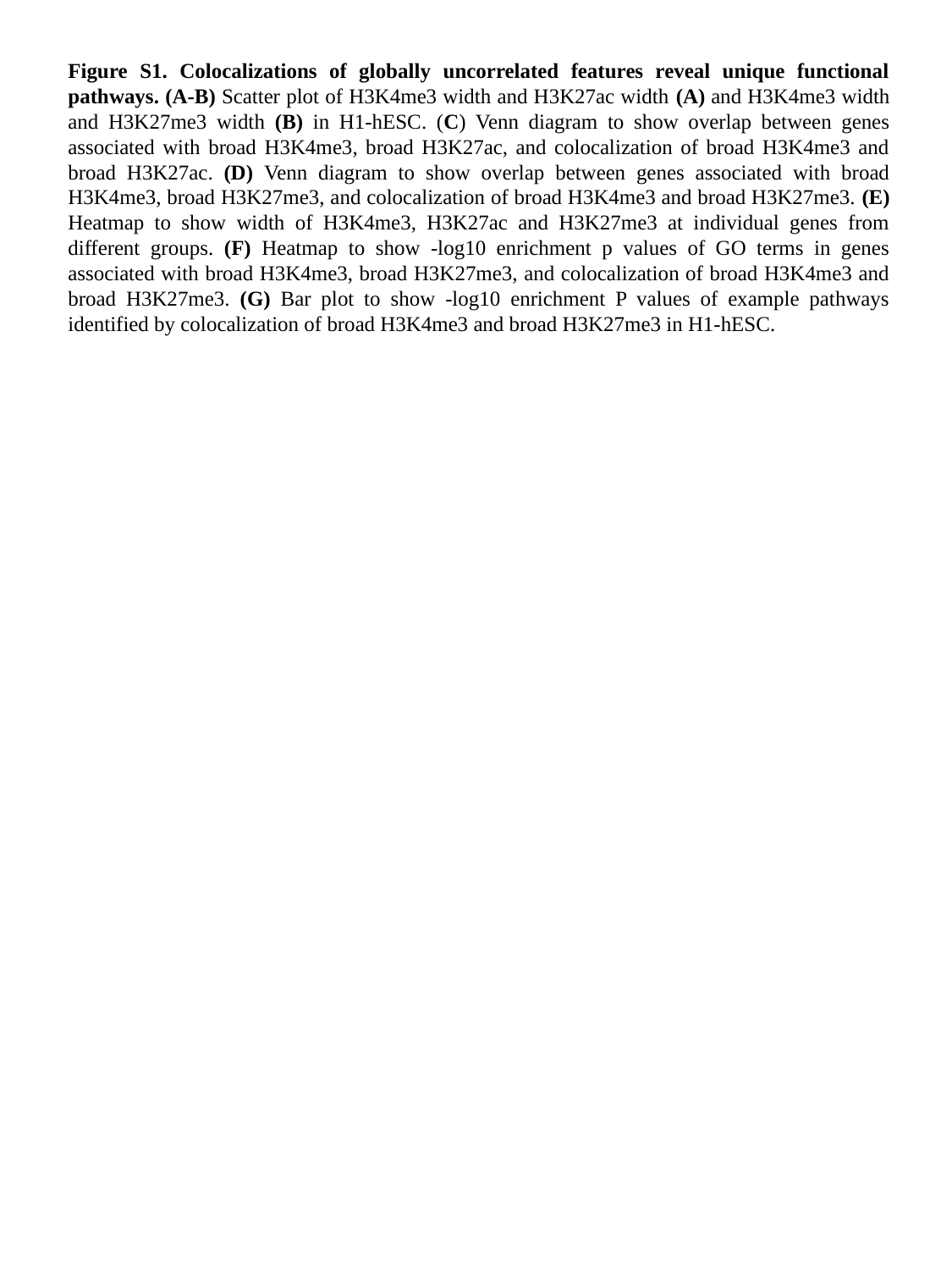

Figure S1. Colocalizations of globally uncorrelated features reveal unique functional pathways. (A-B) Scatter plot of H3K4me3 width and H3K27ac width (A) and H3K4me3 width and H3K27me3 width (B) in H1-hESC. (C) Venn diagram to show overlap between genes associated with broad H3K4me3, broad H3K27ac, and colocalization of broad H3K4me3 and broad H3K27ac. (D) Venn diagram to show overlap between genes associated with broad H3K4me3, broad H3K27me3, and colocalization of broad H3K4me3 and broad H3K27me3. (E) Heatmap to show width of H3K4me3, H3K27ac and H3K27me3 at individual genes from different groups. (F) Heatmap to show -log10 enrichment p values of GO terms in genes associated with broad H3K4me3, broad H3K27me3, and colocalization of broad H3K4me3 and broad H3K27me3. (G) Bar plot to show -log10 enrichment P values of example pathways identified by colocalization of broad H3K4me3 and broad H3K27me3 in H1-hESC.

### Slide 3
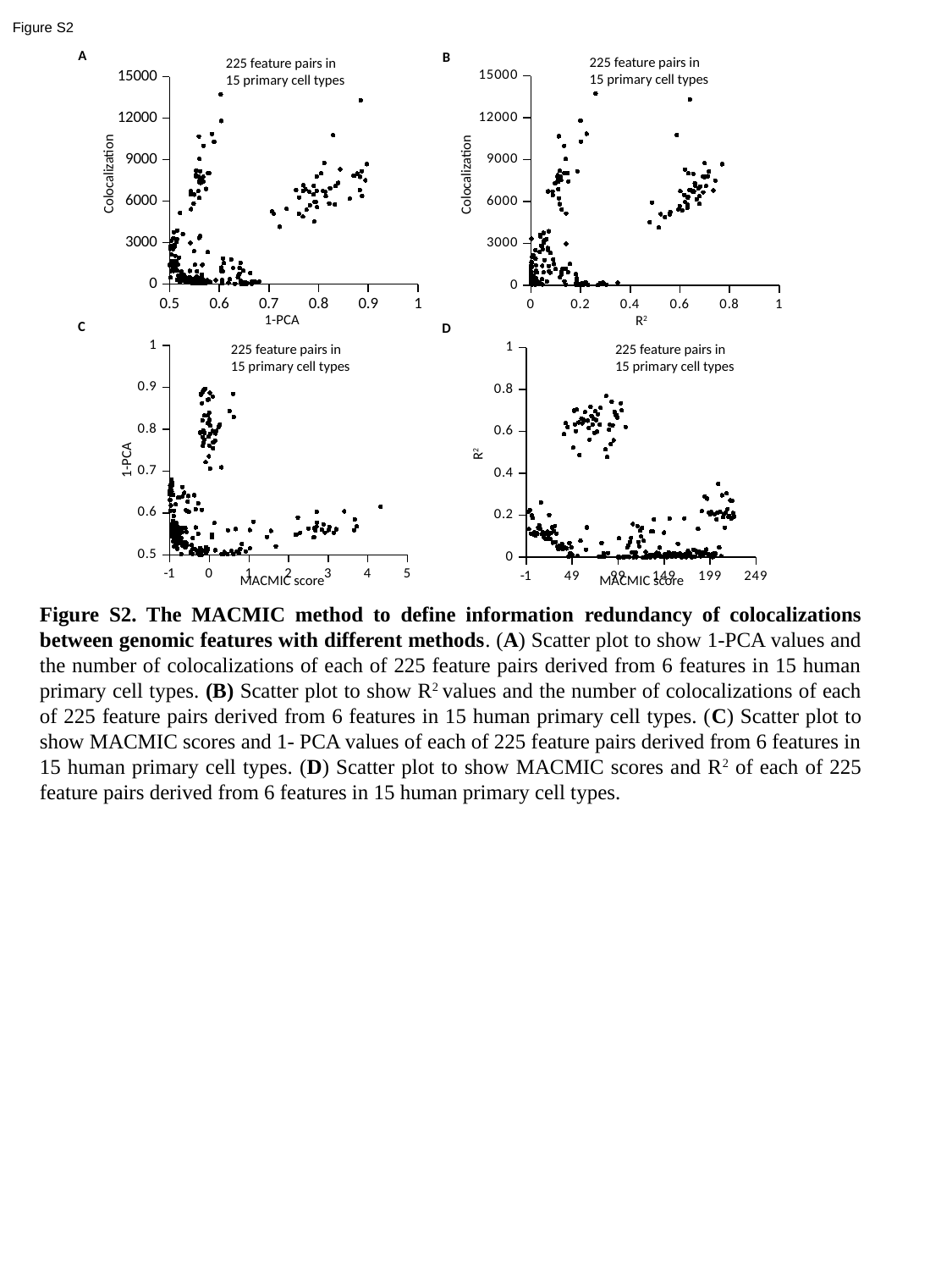

Figure S2
A
B
225 feature pairs in
15 primary cell types
225 feature pairs in
15 primary cell types
#### Chart
| Category | No.overlap.1 |
|---|---|
#### Chart
| Category | No.overlap.1 |
|---|---|Colocalization
Colocalization
1-PCA
R^2^
C
D
#### Chart
| Category | 1-PCA |
|---|---|225 feature pairs in
15 primary cell types
225 feature pairs in
15 primary cell types
#### Chart
| Category | correlation |
|---|---|R^2^
1-PCA
MACMIC score
MACMIC score
Figure S2. The MACMIC method to define information redundancy of colocalizations between genomic features with different methods. (A) Scatter plot to show 1-PCA values and the number of colocalizations of each of 225 feature pairs derived from 6 features in 15 human primary cell types. (B) Scatter plot to show R^2^ values and the number of colocalizations of each of 225 feature pairs derived from 6 features in 15 human primary cell types. (C) Scatter plot to show MACMIC scores and 1- PCA values of each of 225 feature pairs derived from 6 features in 15 human primary cell types. (D) Scatter plot to show MACMIC scores and R^2^ of each of 225 feature pairs derived from 6 features in 15 human primary cell types.

### Slide 4
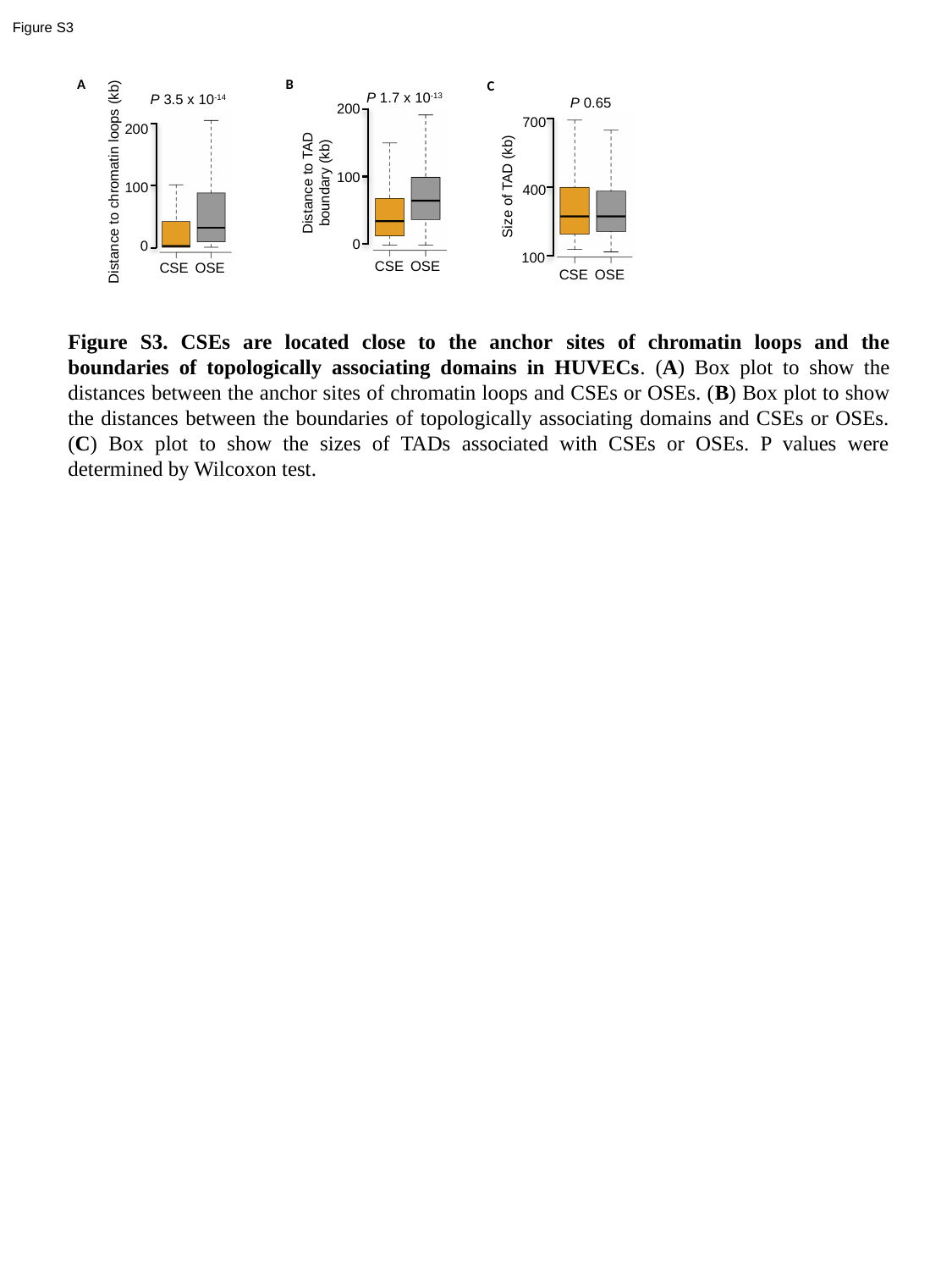

Figure S3
P 1.7 x 10^-13^
P 3.5 x 10^-14^
P 0.65
200
100
0
700
200
100
0
Distance to TAD boundary (kb)
400
Distance to chromatin loops (kb)
Size of TAD (kb)
100
CSE
OSE
CSE
OSE
CSE
OSE
A
B
C
Figure S3. CSEs are located close to the anchor sites of chromatin loops and the boundaries of topologically associating domains in HUVECs. (A) Box plot to show the distances between the anchor sites of chromatin loops and CSEs or OSEs. (B) Box plot to show the distances between the boundaries of topologically associating domains and CSEs or OSEs. (C) Box plot to show the sizes of TADs associated with CSEs or OSEs. P values were determined by Wilcoxon test.

### Slide 5
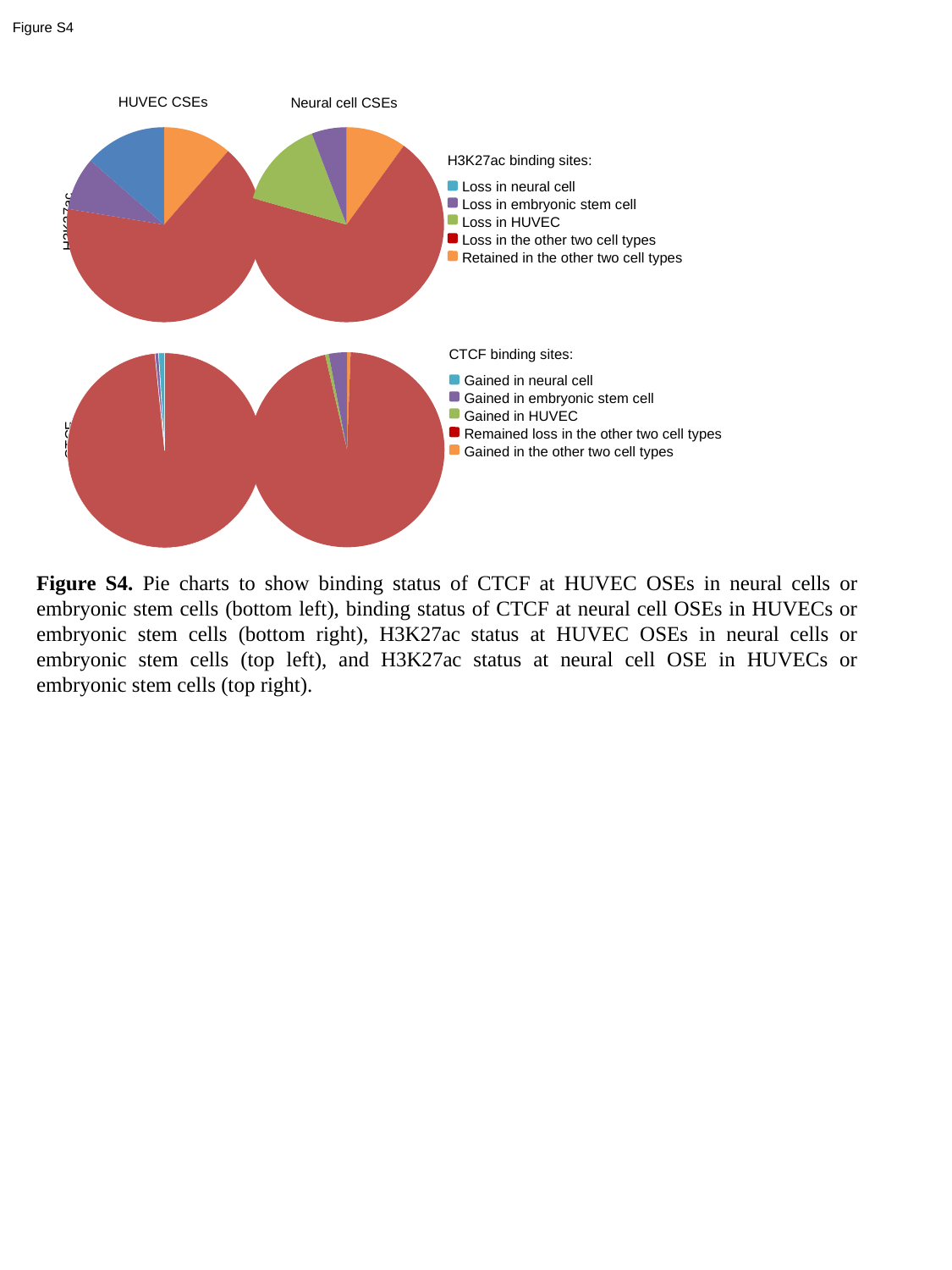

Figure S4
HUVEC CSEs
Neural cell CSEs
#### Chart
| Category | HUVEC |
|---|---|
| retained in both | 57.0 |
| loss in both | 331.0 |
| loss in HUVEC | None |
| loss in embryonic stem | 44.0 |
| loss in neural cell | 68.0 |
#### Chart
| Category | Neural cell |
|---|---|
| retained in both | 50.0 |
| loss in both | 347.0 |
| loss in HUVEC | 74.0 |
| loss in embryonic stem | 29.0 |
| loss in neural cell | None |H3K27ac binding sites:
Loss in neural cell
Loss in embryonic stem cell
Loss in HUVEC
Loss in the other two cell types
Retained in the other two cell types
H3K27ac
CTCF binding sites:
#### Chart
| Category | Neural cell |
|---|---|
| gained in both | 3.0 |
| loss in three | 476.0 |
| gained in HUVEC | 3.0 |
| gained in embryonic stem | 15.0 |
| gained in neural cell | None |
#### Chart
| Category | HUVEC |
|---|---|
| gained in both | 0.0 |
| loss in three | 492.0 |
| gained in HUVEC | None |
| gained in embryonic stem | 3.0 |
| gained in neural cell | 5.0 |Gained in neural cell
Gained in embryonic stem cell
Gained in HUVEC
Remained loss in the other two cell types
Gained in the other two cell types
CTCF
Figure S4. Pie charts to show binding status of CTCF at HUVEC OSEs in neural cells or embryonic stem cells (bottom left), binding status of CTCF at neural cell OSEs in HUVECs or embryonic stem cells (bottom right), H3K27ac status at HUVEC OSEs in neural cells or embryonic stem cells (top left), and H3K27ac status at neural cell OSE in HUVECs or embryonic stem cells (top right).

### Slide 6
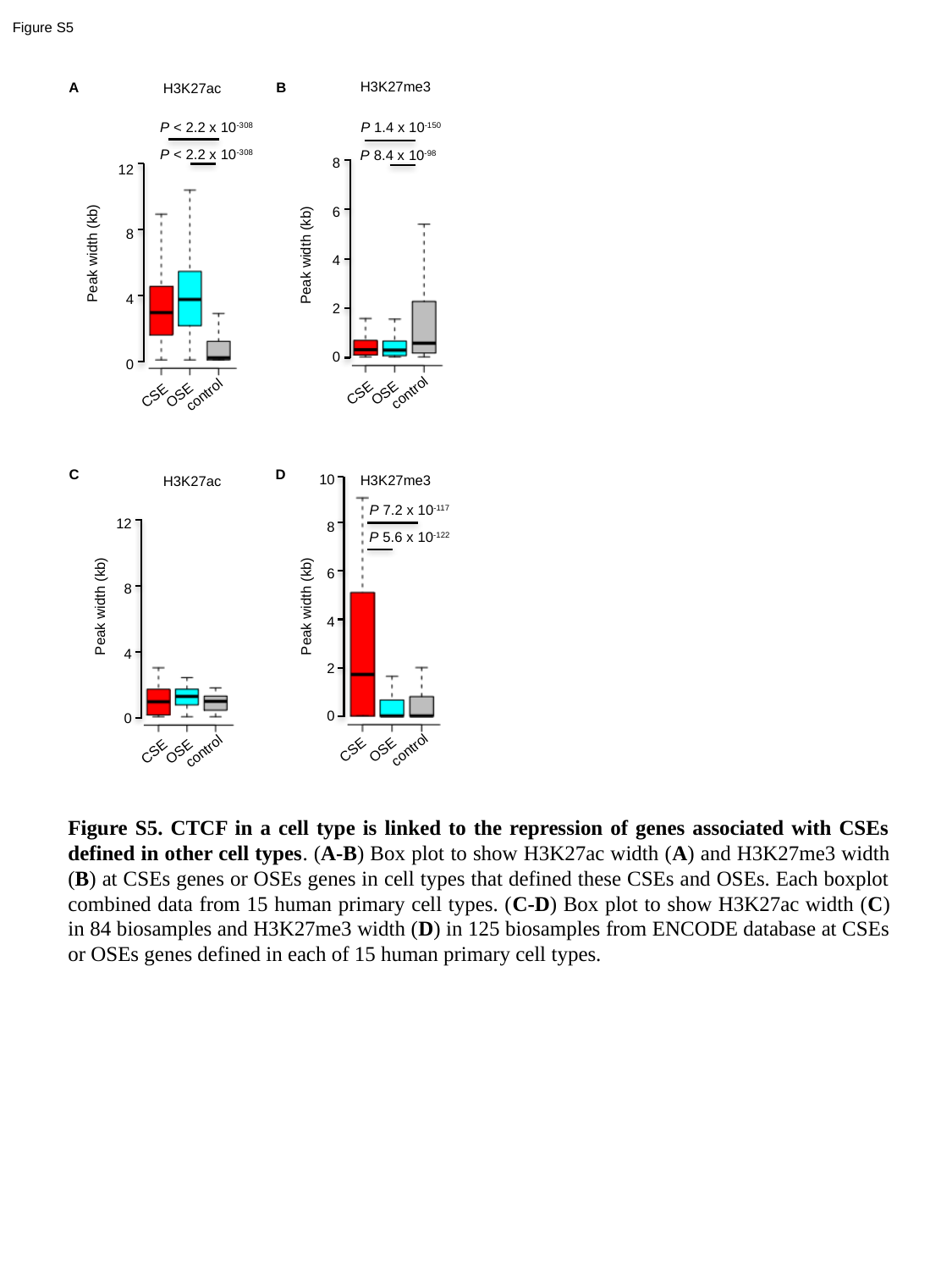

Figure S5
H3K27me3
P 1.4 x 10^-150^
P 8.4 x 10^-98^
8
6
Peak width (kb)
4
2
0
control
OSE
A
B
H3K27ac
P < 2.2 x 10^-308^
P < 2.2 x 10^-308^
12
8
Peak width (kb)
4
0
control
OSE
CSE
CSE
C
D
10
8
6
4
2
0
P 7.2 x 10^-117^
P 5.6 x 10^-122^
Peak width (kb)
Peak width (kb)
control
OSE
control
OSE
CSE
CSE
H3K27me3
H3K27ac
12
8
4
0
Figure S5. CTCF in a cell type is linked to the repression of genes associated with CSEs defined in other cell types. (A-B) Box plot to show H3K27ac width (A) and H3K27me3 width (B) at CSEs genes or OSEs genes in cell types that defined these CSEs and OSEs. Each boxplot combined data from 15 human primary cell types. (C-D) Box plot to show H3K27ac width (C) in 84 biosamples and H3K27me3 width (D) in 125 biosamples from ENCODE database at CSEs or OSEs genes defined in each of 15 human primary cell types.

### Slide 7
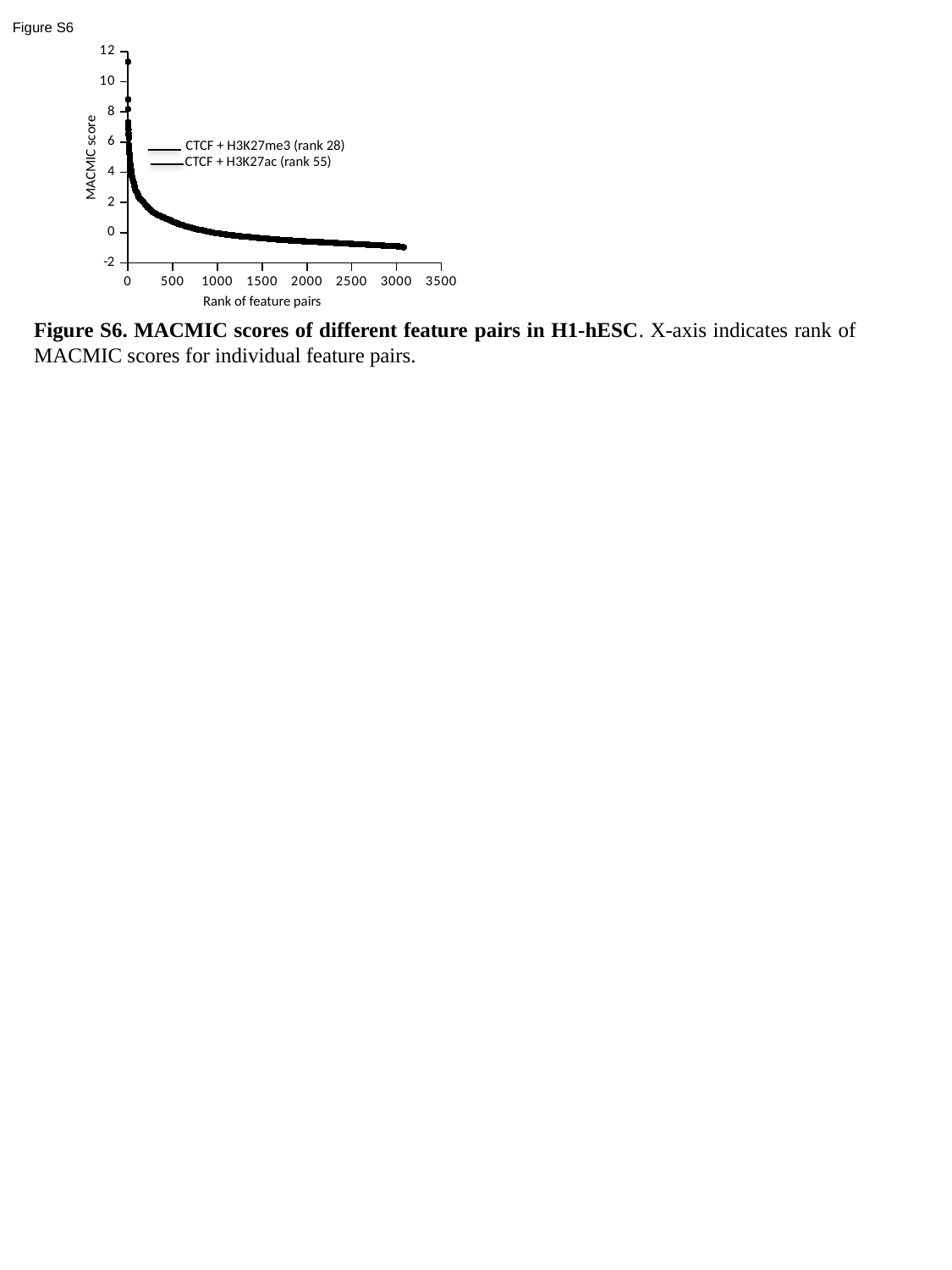

Figure S6
#### Chart
| Category | MACMIC_MI |
|---|---|CTCF + H3K27me3 (rank 28)
MACMIC score
CTCF + H3K27ac (rank 55)
Rank of feature pairs
Figure S6. MACMIC scores of different feature pairs in H1-hESC. X-axis indicates rank of MACMIC scores for individual feature pairs.

### Slide 8
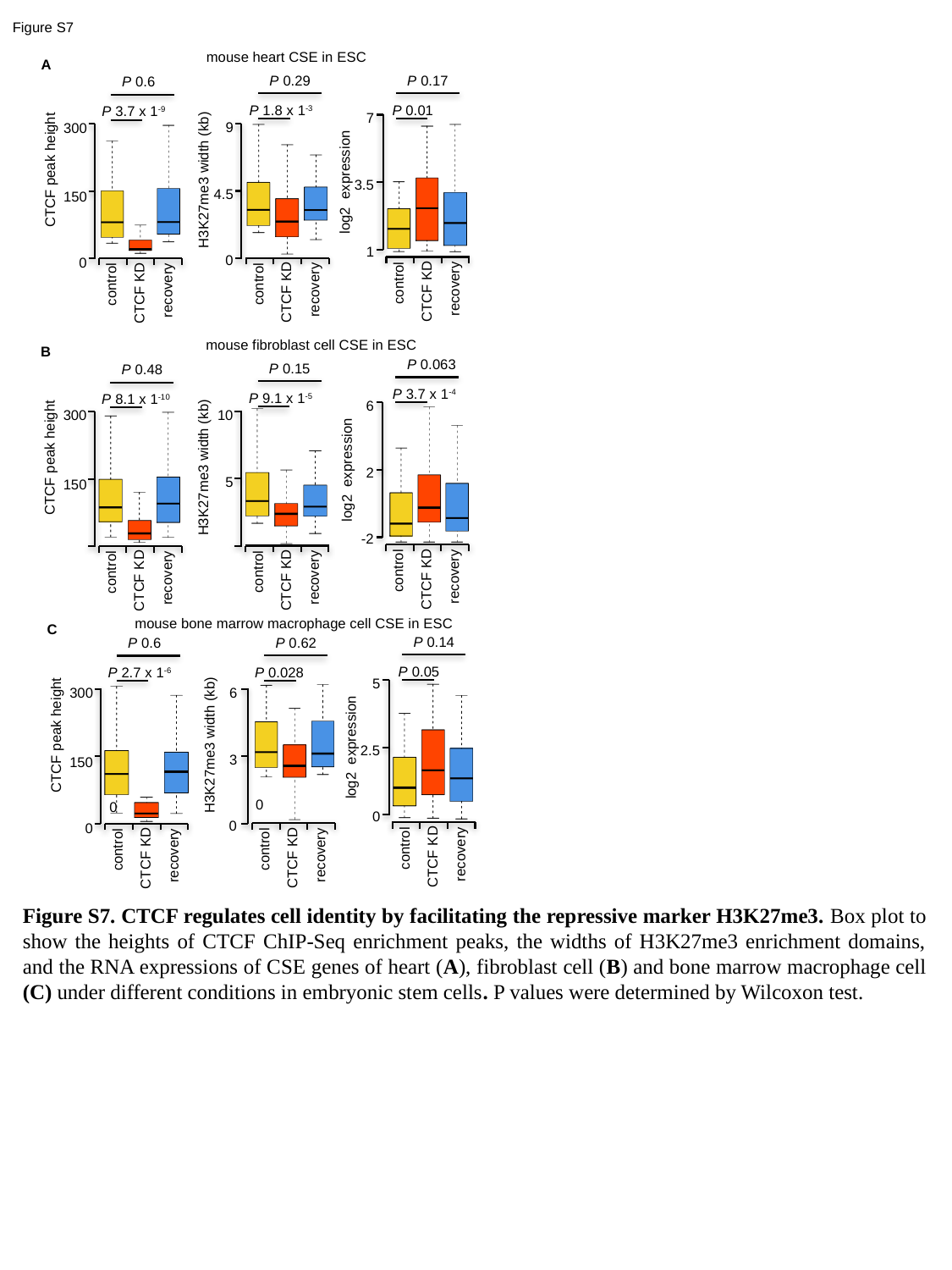

Figure S7
mouse heart CSE in ESC
A
P 0.29
P 1.8 x 1^-3^
P 0.17
P 0.01
P 0.6
P 3.7 x 1^-9^
7
9
300
CTCF peak height
3.5
H3K27me3 width (kb)
log2 expression
4.5
150
1
0
0
control
recovery
CTCF KD
control
recovery
CTCF KD
control
recovery
CTCF KD
mouse fibroblast cell CSE in ESC
B
P 0.063
P 3.7 x 1^-4^
P 0.15
P 9.1 x 1^-5^
P 0.48
P 8.1 x 1^-10^
6
10
300
CTCF peak height
2
H3K27me3 width (kb)
log2 expression
5
150
-2
control
recovery
CTCF KD
control
recovery
CTCF KD
control
recovery
CTCF KD
mouse bone marrow macrophage cell CSE in ESC
C
P 0.14
P 0.05
P 0.62
P 0.028
P 0.6
P 2.7 x 1^-6^
5
6
300
CTCF peak height
2.5
H3K27me3 width (kb)
log2 expression
3
150
0
0
0
0
0
control
recovery
CTCF KD
control
recovery
CTCF KD
control
recovery
CTCF KD
Figure S7. CTCF regulates cell identity by facilitating the repressive marker H3K27me3. Box plot to show the heights of CTCF ChIP-Seq enrichment peaks, the widths of H3K27me3 enrichment domains, and the RNA expressions of CSE genes of heart (A), fibroblast cell (B) and bone marrow macrophage cell (C) under different conditions in embryonic stem cells. P values were determined by Wilcoxon test.
